# DNA sequence-directed cooperation between nucleoid-associated proteins

**DOI:** 10.1101/2020.06.14.150516

**Authors:** Aleksandre Japaridze, Wayne Yang, Cees Dekker, William Nasser, Georgi Muskhelishvili

**Affiliations:** Department of Bionanoscience, Kavli Institute of Nanoscience Delft, Delft University of Technology, Van der Maasweg 9, 2629 HZ, Delft, The Netherlands; Université de Lyon, INSA Lyon, Université Claude Bernard Lyon 1, CNRS UMR5240, Laboratoire de Microbiologie, Adaptation et Pathogénie, 69621 Villeurbanne, France; School of Natural Sciences, Agricultural University of Georgia, Davit Aghmashenebeli Alley 240, 0159 Tbilisi, Georgia

**Keywords:** DNA, Nucleoid Associated Proteins, Atomic Force Microscopy, Nanopores, FIS, H-NS

## Abstract

Nucleoid associated proteins (NAPs) are a class of highly abundant DNA binding proteins in bacteria and archaea. While the composition and relative abundance of the NAPs change during the bacterial growth cycle, surprisingly little is known about their crosstalk in mutually binding to the bacterial chromosome and stabilising higher-order nucleoprotein complexes. Here, we use atomic force microscopy and solid-state nanopores to investigate long-range nucleoprotein structures formed by the binding of two major NAPs, FIS and H-NS, to DNA molecules with distinct binding-site arrangements. We find that spatial organisation of the protein binding sites can govern the higher-order architecture of the nucleoprotein complexes. Based on sequence arrangement the complexes differed in their global shape and compaction, as well as the extent of FIS and H-NS binding. Our observations highlight the important role the DNA sequence plays in driving structural differentiations within the bacterial chromosome.

## Introduction

Nucleoid-associated proteins (NAPs) represent a small class of highly abundant DNA architectural proteins involved in shaping the chromatin as well as in regulating the gene expression in prokaryotes and archaea (1–5). During the bacterial growth cycle, these proteins are expressed in a growth-phase-dependent manner to coordinate the chromosome structure with the metabolic state (6–9). NAPs bind DNA with varying affinities from nanomolar to micromolar concentrations and affect the gene expression by acting as bona fide transcription factors as well as so-called “topological homeostats” (10, 11). The regulation of genomic transcription by NAPs is closely coupled to the availability of free or “unconstrained” DNA superhelicity, as NAPs constrain the DNA acting both as supercoil repositories and topological barriers to supercoil diffusion (11–15).

Factor for Inversion Stimulation (FIS) protein is the most abundant NAP during the exponential growth phase in *Escherichia coli,* while its concentration quickly drops to undetectable levels upon the transition of cells to stationary phase (6, 16, 17). FIS has a global nucleoid-structuring function (18–21) as well as local accessory roles in the assembly of synaptic complexes by site-specific recombinases (22, 23) and transcription-initiation complexes at various promoters, including the exceptionally strong RNA (rRNA and tRNA) promoters (24, 25). The latter are characterized by upstream activating sequences (UAS) containing multiple FIS-binding sites that are arranged in a helical register (26). FIS is also a helix-turn-helix (HTH) DNA-bending protein (27) which upon binding at the phased sites in UAS forms a coherently bent DNA loop that associates with RNA polymerase (28). In general, FIS nucleoprotein complexes form loops and stabilise branches in supercoiled DNA (10, 29–31).

In contrast to FIS, the Histone-like Nucleoid-Structuring (H-NS) protein is a NAP expressed throughout the entire bacterial growth cycle (6), slightly increasing in concentration towards the later growth stages. While overproduction of H-NS *in vivo* is lethal for the cell (32), the deletion of the *hns* gene does not result in large scale restructuring of the nucleoid (18, 33, 34). H-NS is a DNA-bridging protein that binds preferentially to A/T-rich DNA sequences (35, 36), in part mediated by an A-T hook motif that interacts with the DNA minor groove (37, 38). It was shown that H-NS can bridge two DNA helices within a rigid filament (39, 40), trapping RNA polymerase (41, 42). It is assumed that binding of H-NS nucleates at high-affinity sites and subsequently spreads along the DNA strands, leading to gene silencing (43–45). While H-NS can polymerize on a single DNA duplex, resulting in its stiffening (46), it is the DNA-bridging mode facilitated by Mg^2+^ ions that has been primarily implicated in transcriptional repression (47–49).

While the structural role of FIS, H-NS and other DNA-binding NAPs has been intensively studied (18–21, 28, 39, 40), surprisingly little is known about putative cooperative binding effects and the architecture of the ensuing long-range DNA structures. From NAP expression patterns (6) it is clear that distinct combinations of these proteins interact with the genomic DNA during the different stages of the cell cycle (9). Indeed, exploration of their cooperative binding effects appears indispensable for understanding gene regulation. Interestingly, previous studies using Atomic Force Microscopy (AFM) showed that cooperative binding of various combinations of NAPs to the linear phage λ-DNA led to regular structures that were quite distinct from those observed with individual proteins (50), while on binding to large supercoiled molecules, NAPs did phase separate, forming domain-like regions (31). Resolving specific higher-order structures formed by cooperative binding of NAPs is challenging, but it can be achieved by using model DNA sequences that contain a few high-affinity binding sites that facilitate the nucleation of long-range nucleoprotein complexes.

Here, we explore whether the binding of a combination of two major bacterial NAPs, FIS and H-NS, leads to the emergence of distinct nucleoprotein structures that are more than the mere sum of those formed by the individual NAPs. To address this question, we employed DNA sequences with various arrangements of the FIS and H-NS binding sites and studied the resulting higher-order nucleoprotein complexes by use of AFM and nanopore experiments. We find that the sequential organisation of the binding sites directs the assembly of peculiar hairpin-like DNA architectures by FIS and H-NS, which are not observed with either FIS or H-NS alone. Furthermore, we find that FIS and H-NS self-organize to separate domains along the DNA molecules when both proteins are present. Our results exemplify the type of structural rearrangements that local DNA regions can undergo during the bacterial growth cycle, highlighting the crucial role of the DNA sequence.

## Results

### DNA sequence directs cooperative binding of FIS and H-NS

To study the combined effects of FIS and H-NS proteins binding to DNA, we used two plasmids that differed in the sequence arrangement of the NAP-binding sites. In one construct, we arranged a FIS-binding sequence UAS and an H-NS-binding sequence NRE in a Head-to-Tail fashion (HT: UAS-NRE-UAS-NRE). Here, UAS is the upstream activating sequence of tyrosil tRNA gene promoter (*tyrT*UAS)(51) and NRE is the negative regulatory element of proV gene (*proV* NRE) from an osmoregulatory operon (52). In the second construct, the same NAP-binding sequences were arranged in a Head-to-Head fashion (HH: UAS-NRE-NRE-UAS). These sequences were inserted into a 2.9kb backbone devoid of any strong FIS of H-NS binding sites (53) (Figure 1.a & e, Materials and Methods). The 397bp-long *tyrT*UAS region contained three specific FIS binding sites (with *K*_*d*_ values ranging between 7.5 nM and 60 nM) arranged in a helical register (24), while the 264 bp long *proV* NRE contained two high-affinity H-NS sites (*K*_*d*_ values between 15 nM and 25 nM) separated by about 10 helical turns (43). The constructs with Head-to-Tail (HT) and Head-to-Head (HH) arrangements have been described in detail in a previous study (53), which showed that the binding of H-NS to various arrangements of high-affinity binding sites led to the formation of distinct long-range plectonemic coiled structures, that differed in their shape, compaction, and capacity to constrain DNA supercoils.

**Figure 1.**
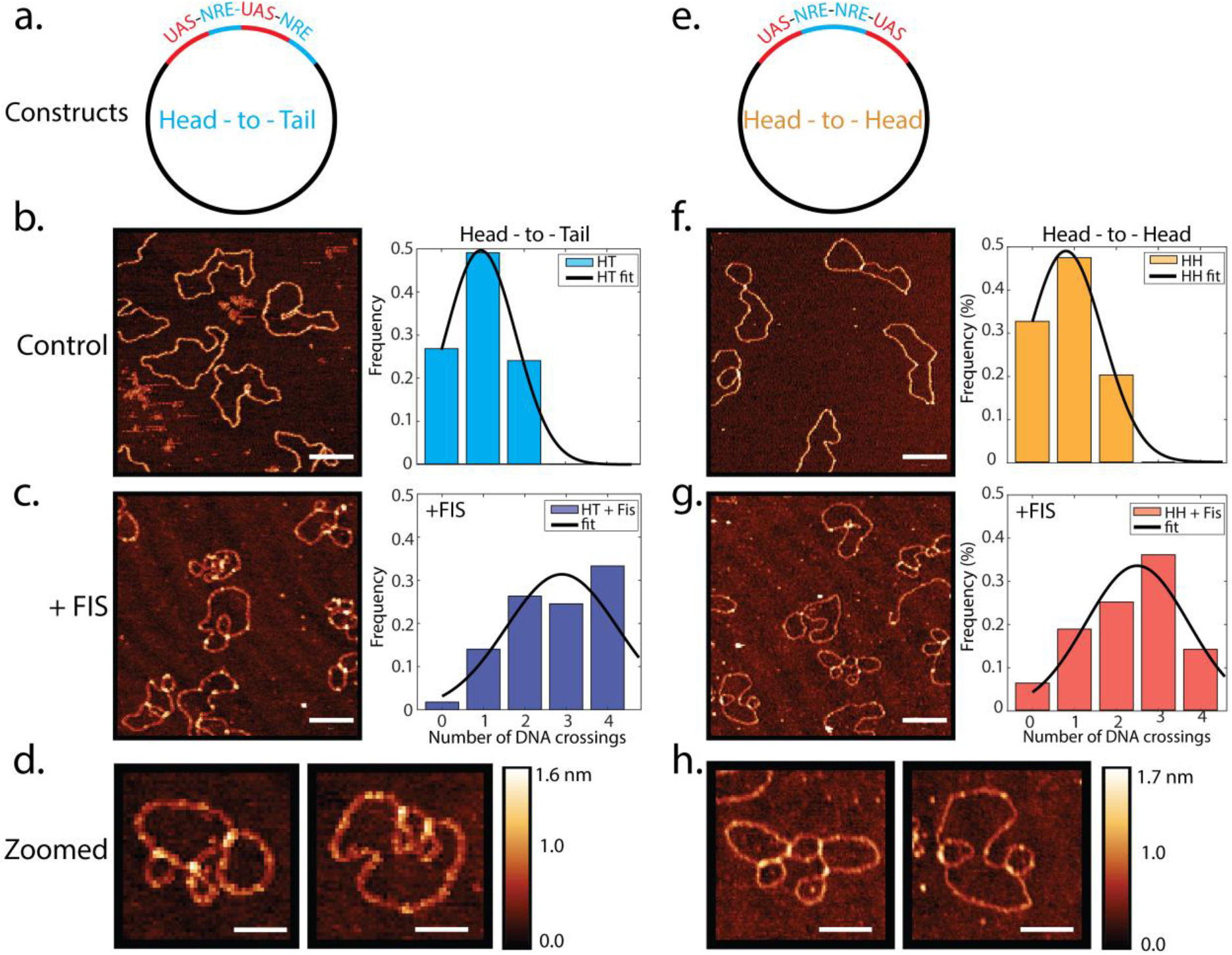
AFM images of FIS nucleoprotein complexes formed on circular DNA constructs. **a.** and **e.** Schematic depiction of the Head-to-Tail (HT) and Head-to-Head (HH) circular constructs. **b.** and **f.** AFM images of control HT and HH plasmids deposited on mica without proteins with the respective number of DNA loops (right panels in b and f). **c.** and **g.** Upon the addition of FIS protein (FIS:bp ratio =0.0026) to the plasmids, the number of DNA loops increased about threefold. Scale bars 200nm. **d.** and **h.** Zoomed AFM images of FIS looped structures. Scale bars 100nm.

First, we investigated nucleoprotein structures formed by the binding of the FIS protein to these constructs. We incubated FIS with either the nicked circular HT or HH plasmid and subsequently imaged by AFM in air (see Materials and Methods). We found that, binding of the protein induced global changes in the shapes of both DNA constructs. FIS stabilised DNA crossovers and loops (Fig.1b-d, f-h). This is in contrast to the plectonemic DNA structures stabilised by H-NS (Supplementary Fig.1). Upon deposition on the surface, naked plasmids displayed, on average, a single DNA crossing (Fig.1b & f, average number of crossings is 1.0 for HT (N=108) and 0.9 for HH (N=129)). Upon co-incubation with FIS, however, the number of loops increased almost three times to 2.8 for HT (N=60) and 2.5 for HH (N=63). Furthermore, the typical height of the DNA crossings stabilized by FIS in both constructs was consistently larger (h_FIS_ =1.35 ± 0.15 nm, N=60; error is sd), than that of DNA crossings in the control samples without protein (h_DNA_=1.05 ± 0.15 nm, N=60; error is sd) (Supplementary Fig. 2). This enabled us to distinguish the FIS binding position along the DNA molecules. Next we looked at the typical loop sizes formed by the binding of FIS protein. The average size of the DNA loops in FIS bound samples was 135 ± 45nm (N=50; error is sd) for the HH construct and 165 ± 45nm for HT (N=50; error is sd), values that in both cases were smaller than the loop size in control sample (230 ± 70nm; N=40; error is sd).

The nanometer resolution of the AFM enabled us to also very precisely measure the various DNA shape parameters, such as the contour length, the radius of gyration, and the effective persistence length of molecules – providing insight into the structural changes induced by the FIS binding (summarized in Table 1). The radius of gyration (R_g_) describes the average globular size of the molecules, while the persistence length (l_p_), describes the stiffness of the DNA molecules. For both constructs binding of FIS decreased the radius of gyration significantly, by roughly 30%, from the initial size of ~150 nm down to ~105 nm (Table 1). Interestingly, when we incubated FIS with a control pBR322 plasmid of a similar size (4.4kb) that was devoid of any strong FIS binding sites (Supplementary Fig. 3), a significantly lower compaction was observed. At similar FIS concentrations, we observed only a 17% compaction. The average number of loops, however, increased by ~3 times (i.e., similar as in the HT and HH constructs) from 0.6 (N=118) to 1.9 (N=85), indicating that this effect was independent of high-affinity FIS binding sites. The effective persistence length of the DNA molecules was also affected by the binding of FIS, decreasing from 52 ± 3 nm (N=35, error is sd) to 40 ± 5 nm (N=30, error is sd) as expected by the observed looping of DNA. Notably, the contour length of the DNA was not much changed by FIS binding (Table 1), in contrast for H-NS binding where we saw a clear compaction.

**Table 1.**
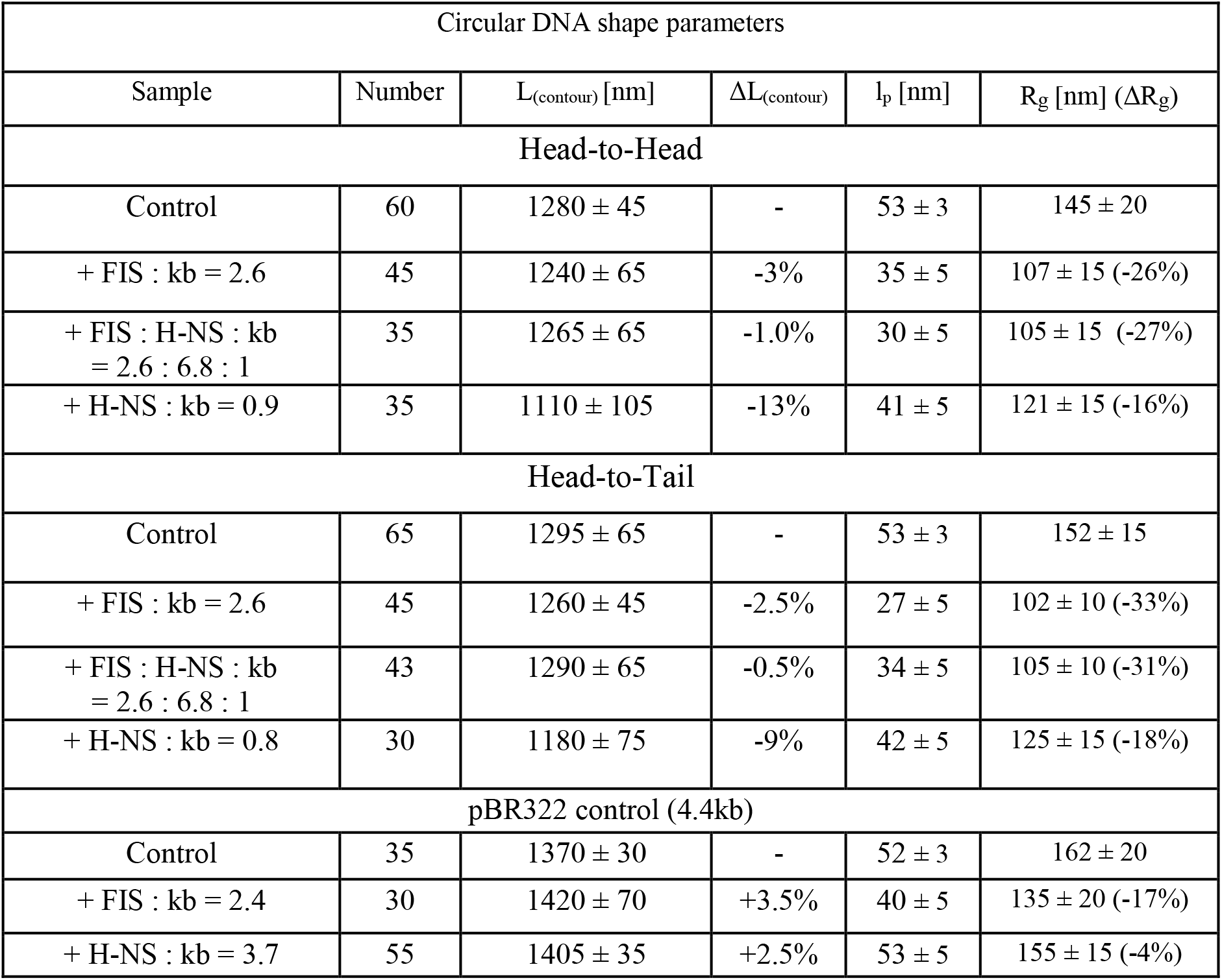
Statistical parameters of DNA molecules bound by FIS and H-NS

Next, we imaged the nucleoprotein complexes formed in the presence of both FIS and H-NS with the HT and HH constructs. In our experiments we used a FIS monomer to DNA bp ratio of 0.0026, and H-NS monomer to DNA bp ratio of 0.0066, protein concentrations that were lower by about an order of magnitude than the maximum physiological levels (6). On both constructs, we observed regions where the DNA duplexes were either bridged into plectonemic braids or were crossing each other to form loops (Figure 2), that is, the characteristic phenomena induced by H-NS and FIS nucleoprotein complexes, respectively. Thus at used near-physiological protein concentrations, both FIS and H-NS could bind naked DNA independently, forming their characteristic nucleoprotein structures. Measurements indicated that the number of DNA crossovers on both constructs was comparable to each other (Figure 2c) as well as to the number of crossovers obtained with FIS alone (Figure 1c and d). By contrast, contour length measurements showed that, on average, the bridged duplex regions formed on HH constructs were significantly more extended compared to HT (Figure 2d). On average, the region with bridged DNA duplexes was 47 ± 2nm in length, (N=43, error is s.e.m.), while the bridged regions observed in HT constructs were much shorter in length, 33 ± 2 nm (N=56, error is s.e.m). Furthermore, on the HH construct, peculiar hairpin-like structures consisting of a “stem” of bridged DNA duplexes with a protruding loop were observed (Fig.2d and e), while no such structures were observed with HT construct, suggesting that they were produced by a cooperative binding effect of FIS and H-NS to a specific (HH) arrangement of cognate binding sites.

**Figure 2.**
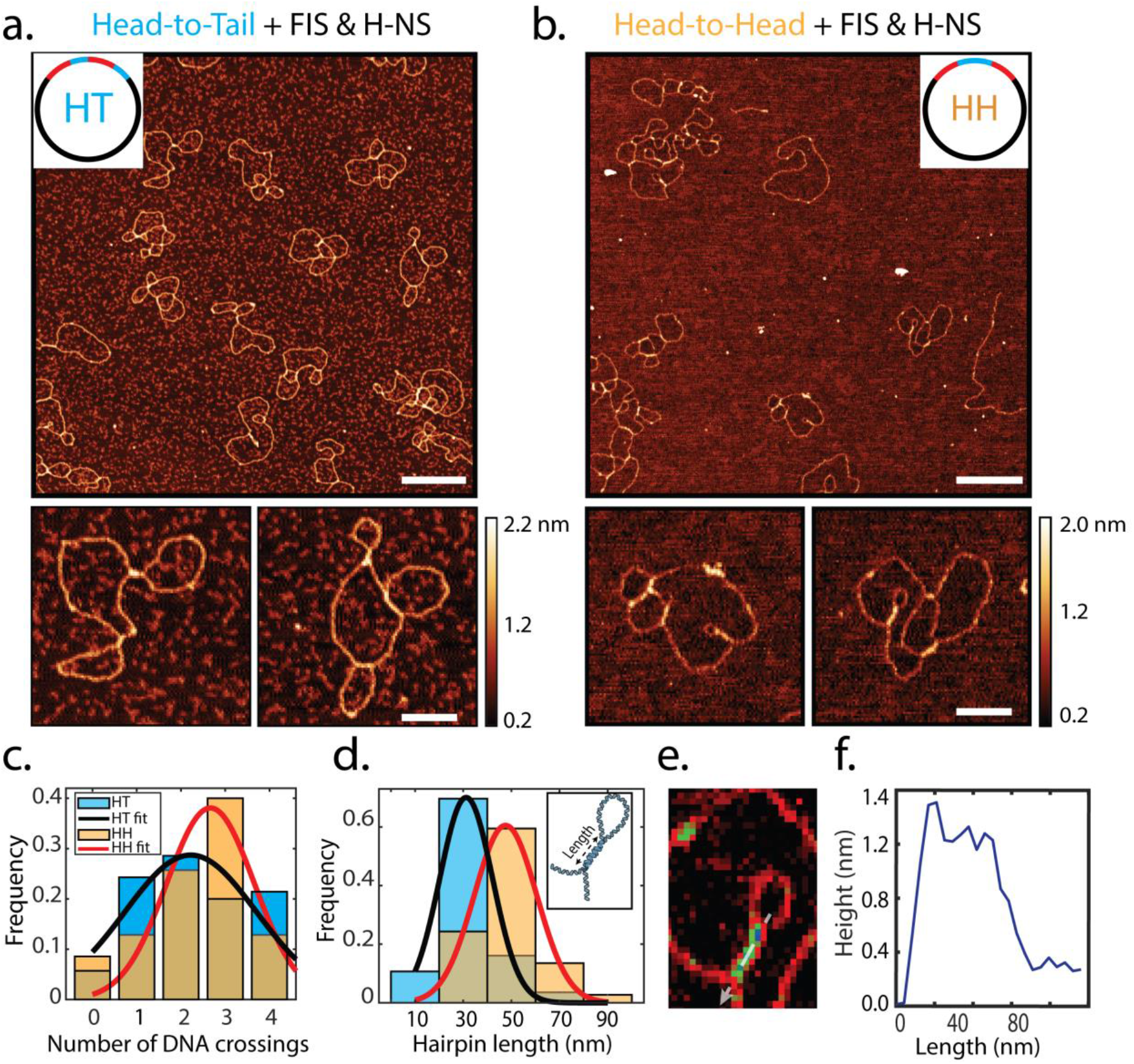
AFM images of FIS and H-NS nucleoprotein complexes formed on HT and HH constructs. **a.** The HT construct forms looped structures indicative of FIS binding, adjoined by DNA bridges indicative of H-NS binding. **b.** The HH construct forms hairpin-like structures where the two proteins appear organised in a specific binding arrangement. Scale bars: 200nm for large scale images and 100nm for zoomed images. **c.** The number of DNA crossings is similar for both HT and HH constructs. **d.** The length of the bridged DNA segments is larger for the HH (N=43) then for the HT (N=56) constructs. Solid lines denote Gaussian fits **e.** Zoomed AFM image of the hairpin structure formed on the HH construct. **f.** Corresponding height cross section (indicated by dotted line on panel e).

Notably, earlier AFM studies measuring the DNA plectonemes stabilised by H-NS on both HH and HT constructs as well as on linear phage Lambda DNA, reported uniform height values of ~ 1 nm along the H-NS bridged regions (50, 53). By contrast, height measurements of H-NS bridged duplexes in the hairpin structures formed on HH constructs with the mixture of FIS and H-NS demonstrated non-uniform heights that varied between the stem of the bridged DNA and the base of the loop, i.e. the point where the DNA duplexes became disjointed (Fig.2,e,f). The stem was typically ~ 1nm in height, while the height of the crossings abutting the diverging duplexes was 1.45 ± 0.2 nm (N=60, error is sd) (Fig.2f), remarkably close to the values obtained for FIS binding (supplementary Fig. 2). Given the propensity of FIS to stabilize DNA loops and high crossovers (28, 30, 31, 40) and that of H-NS to form bridged DNA duplexes of uniform height, it is reasonable to assume that the peculiar DNA hairpins observed in HH constructs represent nucleoprotein complexes formed by cooperative binding of H-NS and FIS at this DNA sequence. We estimate that FIS stabilises the base of the loop whereas H-NS bridges the DNA duplexes and forms the stem of the hairpin. We emphasize that the HT and HH constructs differ only in the spatial arrangement of the sequences with the H-NS and FIS binding sites. Therefore, it is remarkable that binding of H-NS and FIS to these two constructs can lead to the formation of such distinct structures only on one (HH) construct but not on the other (HT).

### Nanopores show that nucleoprotein complexes are different in bulk

To rule out that the differences observed between the HT and HH nucleoprotein complexes represent an artefact of AFM imaging on the mica surface, we investigated the formation of these nucleoprotein complexes in solution by use of solid-state nanopores. Next to serving as a complementary technique to AFM in assessing the formation of nucleoprotein complexes in solution, nanopores also enabled us to gather good statistics (hundreds of DNA translocation events for each experiment). Essentially, the method involves measuring the electrophoretically driven translocation of DNA molecules across a nanometer-sized pore (15 nm diameter in our case) with an applied voltage (see Materials and Methods), as shown in Figure 3a. During the translocation of the molecules across the pore, the DNA temporarily disrupts the flow of ions which leads to a current blockade. As schematically illustrated in Figure 3b, one can distinguish the level of folding and compaction of DNA molecules based on their current blockade levels (54, 55), as well as see if the molecules are bound by proteins (56).

**Figure 3.**
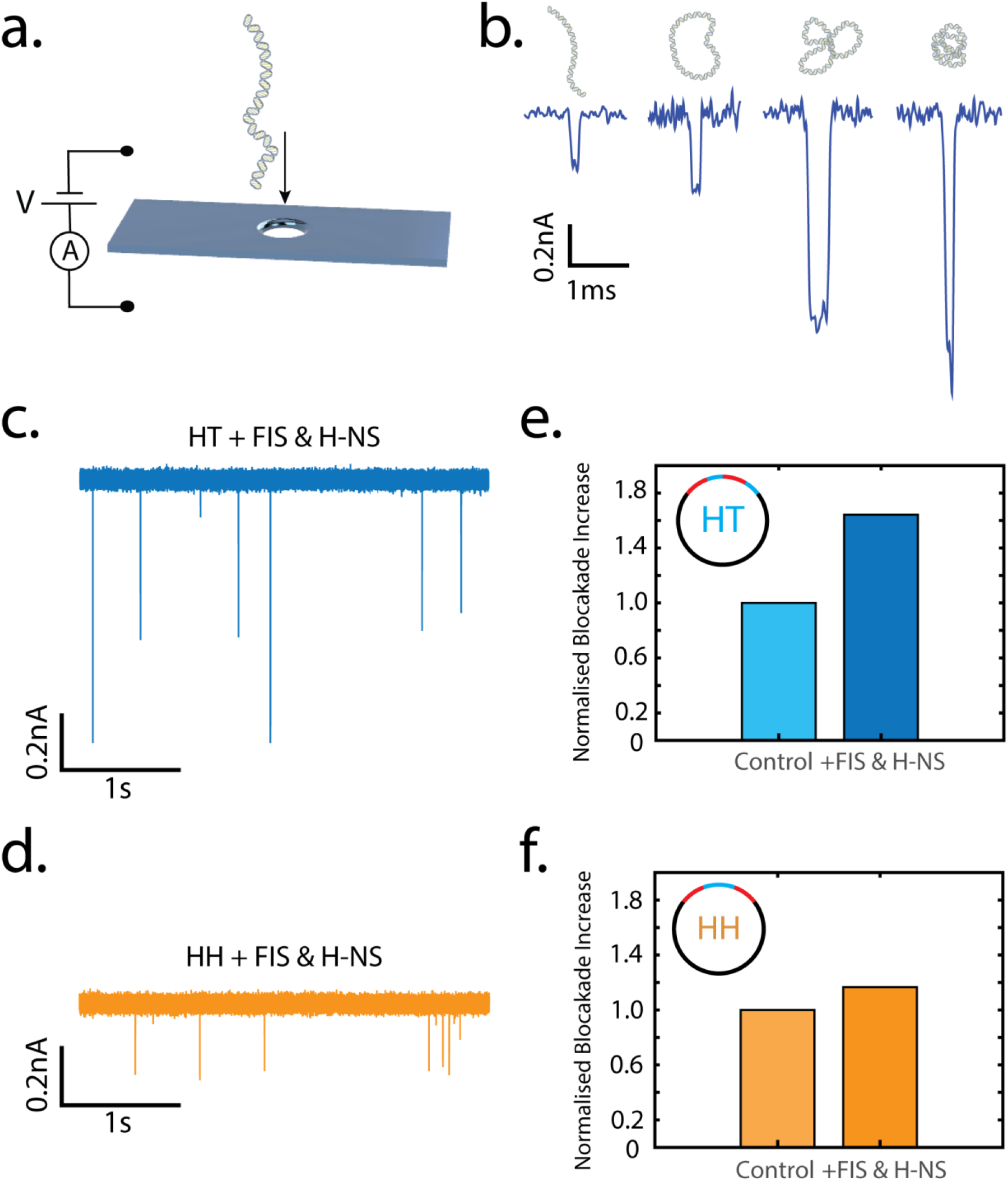
Nanopore experiments. **a.** Schematics depicting the translocation of a DNA through a nanopore. **b.** Example blockade events of HT DNA construct passing through a nanopore. **c.** Typical nanopore traces for the HT + FIS & H-NS samples (1ng/μl final DNA concentration). **d.** Typical nanopore traces for HH + FIS & H-NS samples, (1ng/μl final DNA concentration). Upon the protein addition, the blockades become deeper, indicative of protein binding. The blockade events for the HH + FIS & H-NS samples are uniform and homogeneous, indicating that the DNA-protein complexes are similarly organized. **e.** and **f**. Normalized blockade events for the HT and HH constructs, showing that the deeper events occur much more frequently for the HT construct (in 38% of total events, N_HT+ Protein_=277 out of 737 total events), but not for the HH construct (in 18% of total events, N_HH+Protein_=329 out of 1812 total events).

If the HH and HT nucleoprotein complexes were different in their level of compaction, one would expect to see different blockade levels between the two constructs. As expected, both the HT and HH-DNA plasmid-only controls yielded consistent DNA events with a regular depth in the current blockades (Supplementary Fig. 4). Upon addition of FIS and H-NS, the HT-DNA showed deeper blockade events (Fig.3c) compared to the HH-DNA (Fig.3d). Furthermore, for HH-DNA, the blockade events in the presence of FIS and H-NS were uniform and homogeneous, indicative of a similar organization of nucleoprotein complexes, whereas addition of individual NAPs to HH construct resulted in heterogeneous blockade levels (Supplementary Fig. 4). This finding is consistent with the formation of regular hairpin structures in the presence of both FIS and H-NS. We set a threshold to select for protein bound molecules and then we quantified the percentage of events as compared to the control. Figure 3e and 3f. show the relative increase in the number of deep events for the HT and HH case in the presence of both proteins. We observed a 1.65 fold increase in the number of deep events for the HT construct, but only a 1.19 fold increase for the HH construct. These data confirm that the NAPs binding of the HT construct indeed results in an increased compaction compared to the HH construct (irrespective of the used threshold, Supplementary Fig. 4k).

These differences in organization of HH and HT nucleoprotein complexes in solution were tested further by probing the H-NS nucleoprotein complexes with BamHI restriction endonuclease that cuts the DNA at the junctions of the UAS and NRE elements (Supplementary Fig. 5). We found that the HH construct exhibited a higher protection rate of the junctions compared to HT, indicating that H-NS polymerized more regularly across the UAS/NRE junctions in the HH construct. Taken together, our results indicate that spatial organisation of the FIS and H-NS binding sites in the DNA is determining the 3D architecture of the nucleoprotein complexes, yielding similar DNA structures in the case of HH but not HT orientation of binding sequences.

### DNA sequence arrangement governs the 3D nucleoprotein structure

The only difference between the circular HH and HT constructs resides in the 1.4kb DNA region comprising different arrangements of the UAS and NRE sequences. We studied these 1.4kb regions on linear DNA fragments to test whether they could drive organisation of different nucleoprotein assemblies, and whether the observed distinct hairpin structures could still form with the HH (UAS-NRE-NRE-UAS) as opposed to the HT (UAS-NRE-UAS-NRE) arrangement of the DNA sequence.

Hence we incubated the 1.4 kb linear HH and HT fragments with either FIS or H-NS, or a mixture of both under conditions similar to those used for the circular substrates. We found that binding of FIS alone stabilised looped structures (Fig. 4, Supplementary Fig. 6c), but we observed clear differences between the two constructs. The HT fragment typically formed single ‘lasso’ type structures, while the HH fragment formed ‘butterfly’ structures with multiple loops (Fig. 4c, Supplementary Fig. 6c). The ‘lasso’ structure is consistent with interaction between the FIS-UAS nucleoprotein complexes formed by binding of FIS at the UAS regions located one at the middle and one at the end of the HT fragment, whereas the ‘butterfly’ structure is consistent with interactions of FIS binding at the UAS regions located at the extremities of the HH fragment. The average number of DNA crossings formed upon FIS binding also increased differently for the two constructs. For the HH fragment, the looping number increased by almost 3 times (from 0.67 (N= 104) to 1.95 crossings (N= 80)), while for the HT it increased by more than twice (from 0.55 (N= 111) to 1.25 crossings (N= 95)) compared to samples without the added protein. Interestingly, for both constructs, FIS formed DNA loops of a similar size ~100nm in contour length (HH L_loop_HH_=110 ± 35 nm (N=43, error is sd) and HT L_loop_HT_=100 ± 25 nm (N=40, error is sd), i.e., smaller in size compared to the loops formed by simple deposition of the naked DNA on surface L_loop_control_=150 ± 60 nm (N=85, error is sd) (Supplementary Fig. 7). Similarly, as in the case of circular DNA molecules, the binding of FIS compacted the structures thereby reducing the radius of gyration as well as the effective persistence length (Supplementary Table 1).

**Figure 4.**
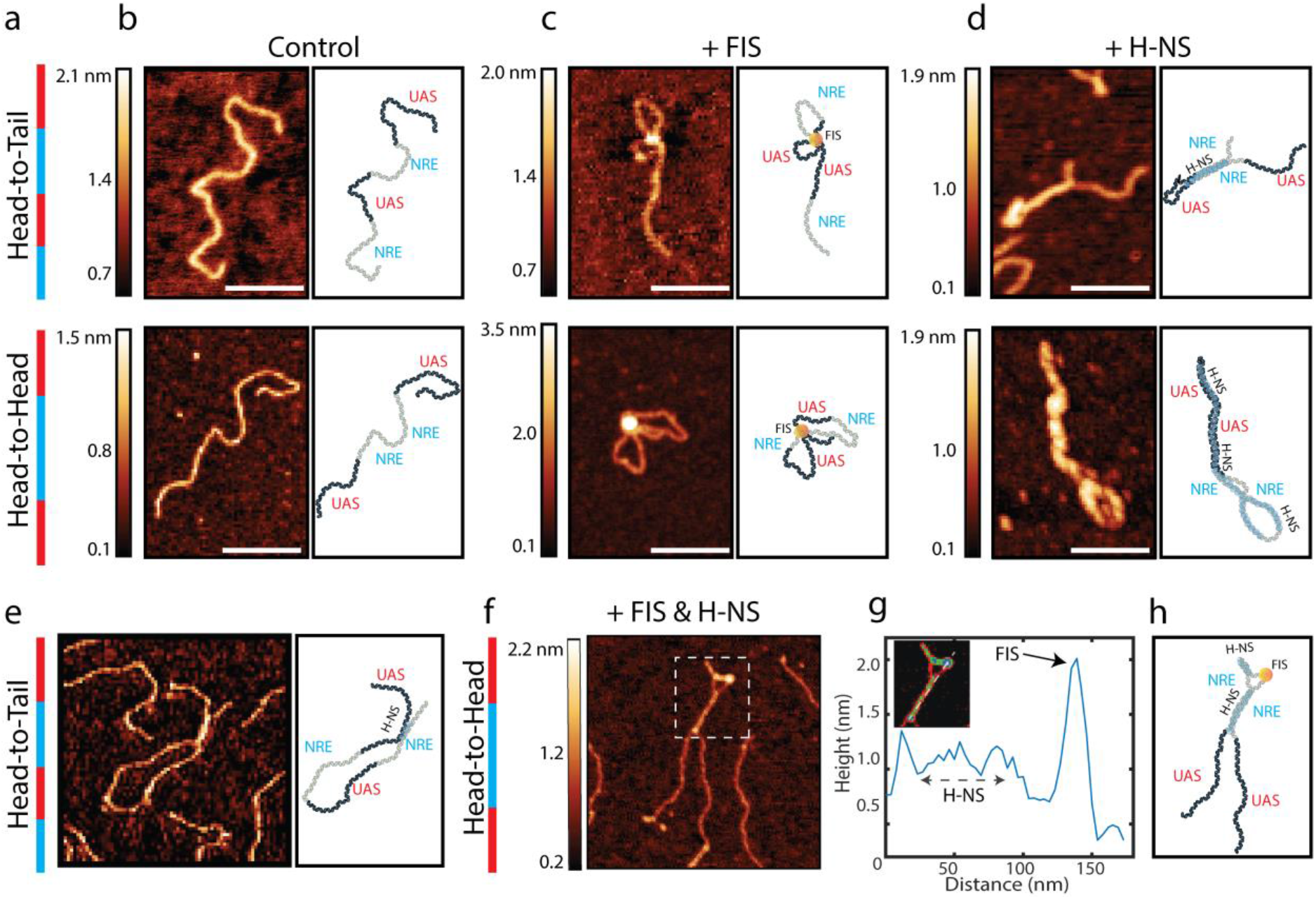
AFM images of nucleoprotein complexes formed on linear 1.4kb fragments. **a.** Schematic depiction of the Head-to-Tail (HT) and Head-to-Head (HH) constructs. Blue indicates the NRE and red UAS regions. **b-h.** AFM images with corresponding schematic depictions of the nucleoprotein organisation for **b.** control, **c.** FIS-bound and **d.** H-NS-bound linear fragments, **e.** nucleoprotein complex formed by HT fragment bound by FIS & H-NS, **f-h.** nucleoprotein complex formed by HH fragment bound by FIS & H-NS. **g.** Height measurement of the nucleoprotein complex formed up on binding of FIS & H-NS to the HH fragment shown on panel f. Scale bars 100nm.

For the H-NS only condition, we also observed different bridged DNA duplex structures on the HT and HH linear fragments (Supplementary Fig. 6d). For the HT fragment, the bridged duplexes were associated with extruding ends of unequal lengths, consistent with H-NS bridging of the NRE sequences located once at the end and once in the middle of the fragment. For the HH fragment, the bridged duplexes did not demonstrate any extruding ends, consistent with a previously observed configuration (53), in which the two adjacent NRE regions engaging all the four consecutive high-affinity nucleation sites are collinearly intertwined (Fig. 4d). Unsurprisingly, the bridged regions were of different lengths, being shorter in HT arrangement, L_bridge_HT_ = 45 ± 25 nm (N=43, error is sd) compared to HH L_bridge_HH_ = 65 ± 40 nm (N=36, error is sd). What was surprising to see, was that on both the HT and HH fragments, the looped DNA regions connecting the H-NS-bridged duplexes appeared to be also covered by H-NS (Supplementary Fig. 8). This demonstrated that under the same environmental conditions H-NS could polymerize on individual DNA duplexes (48) as well as form bridges between two DNA strands (31, 39, 50, 53) based on the DNA sequence alone.

When a mixture of FIS and H-NS was incubated with linear HT fragments, no regular structures were observed, as was the case with circular HT DNA (Fig. 4e). Strikingly, with HH fragments however, the binding of FIS and H-NS again formed hairpin-like structures consisting of a stem of bridged DNA duplexes associated with a loop, resembling those formed with circular HH DNA (Fig. 4f, Fig. 5). Measurements revealed uniform heights across the H-NS bridged regions, whereas blobs of greater heights were associated with the loops, indicative of FIS binding. We infer that the differences observed between the structural organization of nucleoprotein complexes formed with HT and HH constructs are driven by distinct spatial arrangements of the UAS and NRE elements containing the FIS and H-NS binding sites (Fig. 4h and Fig.5).

**Figure 5.**
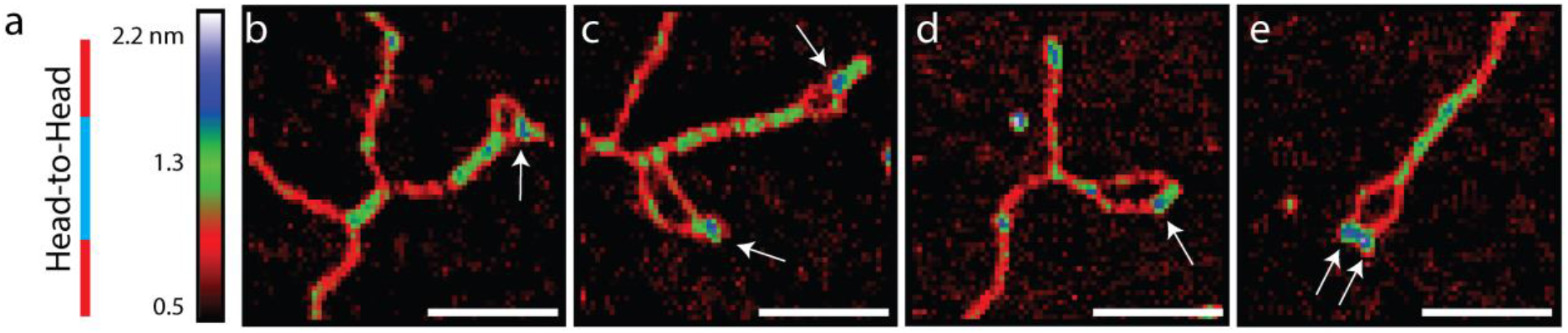
FIS and H-NS proteins phase separate when binding to the DNA. **a.** Schematic depiction of the Head-to-Head (HH) construct. Blue indicates the NRE and red UAS regions. **b.-e.** Zoomed AFM images of HH fragments with simultaneously bound H-NS and FIS proteins. White arrows indicate the position of FIS proteins. Scale bars 100nm.

## Discussion

In this study we set out to investigate the role of DNA sequence organisation in the assembly of higher-order nucleoprotein structures formed by simultaneous binding of two abundant bacterial NAPs, FIS and H-NS, to different spatial arrangements of their cognate DNA binding sites.

We chose FIS and H-NS because of their competing functions *in vivo*. The global “genomic silencer” H-NS forms tightly interwound plectonemic structures which impede transcription (2, 41, 49), whereas open DNA structures, such as toroids and DNA loops are stabilised by the global transcriptional activator FIS, are conducive to transcription initiation (28–30). Furthermore, both FIS and H-NS were found to be simultaneously present in transcriptionally active regions *in vivo* (57), where they appear to compete with each other for binding by stabilising different DNA topologies (47). Importantly, while some major factors involved in regulating the global architecture of the bacterial genome become progressively resolved (18, 19, 34, 58), the smaller-scale local architectures remain largely obscure. Given the large variations in the NAP protein composition during the growth cycle of the cell (6), it is fair to assume that the nucleoid undergoes reshaping (59) with corresponding changes in the local architecture depending on the metabolic state of the cell.

To study the nucleoprotein complexes formed by combination of FIS and H-NS, we used two DNA substrates with a spatially different, Head-to-Head or Head-to-Tail arrangement of the NAP binding sites. Binding of individual NAPs to the circular plasmids demonstrated a conspicuous difference between the architectures of the FIS and H-NS nucleoprotein complexes, fully consistent with previous microscopic observations using supercoiled DNA (31, 40). Interestingly, in contrast to H-NS, we did not find any clear differences between nucleoprotein complexes formed on the two substrates with FIS alone. However, when a combination of FIS and H-NS was used, we found that various arrangements led to structurally distinct nucleoprotein complexes. In particular, binding of FIS and H-NS to the HH arrangement of sites produced specific hairpin-like structures in which the bridged duplexes stabilised by H-NS were associated with a loop apparently stabilised by binding of FIS, as indicated by the AFM height measurements (Fig. 2). This observations is consistent with earlier studies showing that FIS can stabilise both DNA loops and crossovers (30, 40) while the competition between FIS and H-NS induces alternative DNA curvatures in the UAS regions (47). When we repeated the experiments with linear DNA fragments containing just the sequences with high-affinity protein binding sites, similar hairpin structures were observed with the HH but not the HT substrate (Fig. 4, Fig. 5). Since we could not observe such regular structures with the HT arrangement of binding sites with either the circular of linear DNA substrates, we infer that their peculiar architecture is a result of crosstalk between FIS and H-NS driven by the specific spatial organisation of the binding sequences.

In nanopore experiments, we found that the HH nucleoprotein complexes formed by FIS and H-NS demonstrated regular current blockade levels in contrast to HT (Fig. 3, Supplementary Fig. 4). Furthermore, restriction endonuclease experiments showed enhanced protection of the NRE-UAS junctions in nucleoprotein complexes formed on HH compared to HT constructs (Supplementary Fig. 5) confirming our previous observations (53). Both these features are consistent with the AFM data and suggest a higher structural regularity of the HH nucleoprotein complexes compared to that of the HT nucleoprotein complexes.

The organisation of HH nucleoprotein complexes was shown to be due to H-NS bridging of the two adjacent NRE elements that contain high-affinity H-NS binding sites and plectonemic coiling of the DNA (53). This configuration leads to extrusion of tightly bent small loops that are bound by FIS, as observed in our experiments (Fig. 2 & 4). Such tight loops might be especially attractive for further FIS binding. First, a tightly bent DNA loop would facilitate FIS binding due to unusually short distance between the helix-turn-helix motifs of FIS and accordingly, strong preference for tightly bent DNA substrates with compressed minor groove (27). Second, while the NREs demonstrate a sequence periodicity of ~11 bp, H-NS-bridging of the adjacent NREs in HH construct constrains negative superhelicity, most likely due to the right-handed plectonemic coiling and stabilisation of high negative twist (53). This would lead to compensatory over-twisting of the extruding loop reducing the helical repeat to values <10.5 bp, consistent with the assumed preferential value for FIS binding of ~10.2 bp (60). This over-twisting would also counteract the H-NS binding within the tight loop stabilised by FIS and lead to organisation of structures observed in Fig.5, in which FIS is positioned in the looped DNA regions surrounded by H-NS bridged DNA filaments. Interestingly, in the absence of FIS, H-NS is capable of polymerising on the tightly bent loop connecting the bridged helices (Fig.4d, Supplementary Fig.8), meaning that FIS hinders the spread of H-NS. Notably, recent studies demonstrated that the physiological B-form DNA can undergo structural changes resulting in non-canonical DNA forms under the influence of DNA supercoiling (61), spatial confinement (62) and DNA sequence (63, 64), indicating that the DNA structure itself is highly heterogeneous.

The local arrangement of FIS and H-NS proteins in the HH hairpins is non-random. Some form of polymer-driven phase separation could drive a differentiation of chromatin architecture (65, 66), in keeping with the reported capacity of FIS and H-NS to form topological barriers in the chromosome (13). Looped regions stabilised by FIS that are expelling H-NS, could potentially be accessible to transcription machinery, creating genomic islands of ‘open’ and ‘closed’ DNA regions. These coexisting structures are in line with the predominantly repressing and activating roles of H-NS and FIS on transcription, respectively. Since the distribution of the H-NS and FIS binding sites in the *E. coli* genome is non-random (36, 67), we hypothesize that the growth-phase-dependent global regulation of transcription (68) involves alterations of chromatin accessibility. This accessibility of chromatin is further modulated by a crosstalk between temporally changing composition of NAPs and a non-random distribution of their cognate genomic binding sites, thus leading to spatiotemporal organisation of the ‘open’ and ‘closed’ DNA structures that are crucial to genomic expression.

## Materials and Methods

### Constructs

The two 3997 bp constructs (Head-to-Tail and Head-to-Head) were constructed as described earlier (53). Briefly, the constructs contained sequences with FIS binding sites amplified from the UAS of the *tyrT* gene (denoted as UAS) and sequences with H-NS binding sites amplified from the NRE of *proV* gene (denoted as NRE) of *E. coli*. In these two constructs the individual U and P elements were cloned in different spatial arrangements. Fis and H-NS were purified as previously described (69, 70).

### Linear DNA sample preparation

Linear DNA fragments were diluted in AFM buffer (20mM Hepes, 50mM KCl, 2 mM NiCl_2_, 0.003% Tween 20, 2.5% Glycerol, pH 7.9) to a final concentration of 10 ng/μl. A control sample, without proteins, was prepared by mixing one microliter of linear fragment DNA with 22 μl of AFM buffer and incubating for 5 min at room temperature on freshly cleaved mica. The mica was then rinsed with 1 ml of ultrapure water and dried under a gentle flow of compressed filtered air.

For protein-DNA constructs first both H-NS and FIS were first diluted in the AFM buffer to the desired concentration (FIS - 80 ng/μl, HNS- 22 ng/μl). Several samples at different protein to DNA ratios were then prepared for each construct.

Protein-DNA samples were prepared by mixing 1 μl of protein dilution with 21 μl AFM Buffer and only after mixing the proteins solution well, 1 μl of template DNA (10 ng/μl) was added to the solution. The whole mix (23μl) was incubated for 5 min at 37 °C. As the pre last step, the incubation mix was deposited on freshly cleaved mica surface and incubated at room temperature for 5 min. Finally, the surface was rinsed with 1ml ultrapure water and dried under gentle nitrogen flow.

### Circular DNA sample preparation

Circular DNA constructs were nicked using the Nt.BspQI nuclease (New England BioLabs) and purified from 1% agarose gel. DNA was then diluted in the P1 buffer (1 mM TRIS-HCl, 4 mM MgCl2, 0.003% Tween 20, 2.5% Glycerol, pH 7.9) to a final concentration of 10 ng/μl. Control samples, without proteins, were prepared by mixing 1 μl of DNA with 22 μl of P1 buffer and incubated for 7 min at room temperature on freshly cleaved mica. The mica was then rinsed with 1 ml of ultrapure water and dried under a gentle flow of compressed filtered air. Protein-DNA samples were prepared by mixing 1 μl of protein dilution (in P1 Buffer) with 21 μl P1 Buffer and only after mixing the proteins solution well, 1 μl of template DNA (10 ng/μl) was added to the solution. The whole mix (23μl) was incubated for 5 min at 37 °C. As the pre last step, the incubation mix was deposited on freshly cleaved mica surface and incubated at room temperature for 5 min. Finally, the surface was rinsed with 1ml ultrapure water and dried under gentle nitrogen flow.

### AFM imaging

Images were collected using a Multimode atomic force microscope equipped with a Nanoscope IIIa controller (Veeco Instruments, Santa Barbara, CA), operating in Tapping Mode in air using a J-scanner and RTESP silicon cantilevers. All recorded AFM images consist of 512×512 pixels with scan frequencies between 1 and 2 Hz. Each protein-DNA binding experiment was performed at least in duplicate. AFM images were obtained at several separate locations across the mica surface to ensure a high degree of reproducibility and were used for statistical analysis of protein-DNA complexes. Only DNA molecules that were completely visible in AFM image were considered for statistical analysis. AFM images were simply flattened using the Gwyddion software (Version 2.55) without further image processing (71).

### Analysis of AFM images

Statistics properties such as the contour length and the radius of gyration of DNA molecules (typically between 35–100 individual molecules) were analysed using “DNA Trace” software (72). We measured the effective persistence length *lp* of control and protein bound DNA molecules by using the bond correlation function for polymers in two-dimensions,

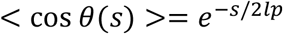

where *θ* is the angle between the tangent vectors to the chain at two points separated by the distance *s*, and *lp*, the persistence length (73). This analysis is a proven method to extract the persistence length for polymer molecules that orient themselves in an unconstrained way at a surface in equilibrium. We apply the approach here as well to nucleoprotein filaments that contain small loops, and hence denote *lp* as the effective persistence length.

### Nanopore experiments

We used TEM-drilled 15 nm diameter SiN nanopores for the experiments. The SiN membrane containing the nanopore was loaded in a PEEK (Polyether ether ketone) flow cell. The DNA samples were diluted in 1M LiCl to a final concentration of 1ng/μl before being introduced to the cis side (−ve) of the nanopore. We used Ag/AgCl electrodes and an Axopatch 200B amplifier (Molecular Devices) for current detection. In experiments where FIS (5.6ng/μl incubation concentration) and H-NS (6ng/μl incubation concentration) were used, the DNA molecules (10ng/μl) were pre-incubated with proteins for 10min at room temperature in 1M LiCl buffer and then diluted 10 times (final DNA contecntration 1ng/μl in into 1M LiCL) added to the cis side of the nanopopore. The traces were recorded at 200 kHz and further low pass filtered at 10 kHz with the Transanalyzer Matlab package (74). As DNA purification quality as well as self-folding of the DNA duplex may affect the current blockade levels, we normalized the blockade levels for each construct to set the control standard for comparison of the protein-bound constructs. We set a current blockade threshold at 2.5 times higher than the average current blockade of a single DNA helix event (threshold = 2.5·I_DNA_) and quantified the percentage of events above this threshold as compared to the control. Variation of the thresholding level had nearly no effect on the results, as can be seen in the Supplementary Fig.4k.

### Probing with restriction enzymes

The cleavage reactions of the FIS & H-NS nucleoprotein complexes were carried out for 5 min with BamHI (New England BioLabs) restriction enzyme for 10 min at 37 °C in the standard New England BioLabs CutSmart buffer. Fis & H-NS were added in increasing concentrations to DNA constructs and incubated for 6 min at 37 °C before the addition of restriction enzymes. The samples containing reaction products were loaded on 1% agarose gel, and the intensities of individual DNA bands were analyzed using Gel analysis option on ImageJ software.

## Acknowledgements

The work was supported by the Netherlands Organisation for Scientific Research (NWO/OCW), as part of the NanoFront and BaSyC programs. C.D. acknowledges support by ERC Advanced Grant SynDiv (no. 669598). A.J. acknowledges support by the Swiss National Science Foundation (Grants P2ELP2_168554 and P300P2_177768).

