## Supplementary data for "DNA sequence-directed cooperation between nucleoid-associated proteins"

**This SI includes:**

Supplementary Figures 1-8

Supplementary Table 1

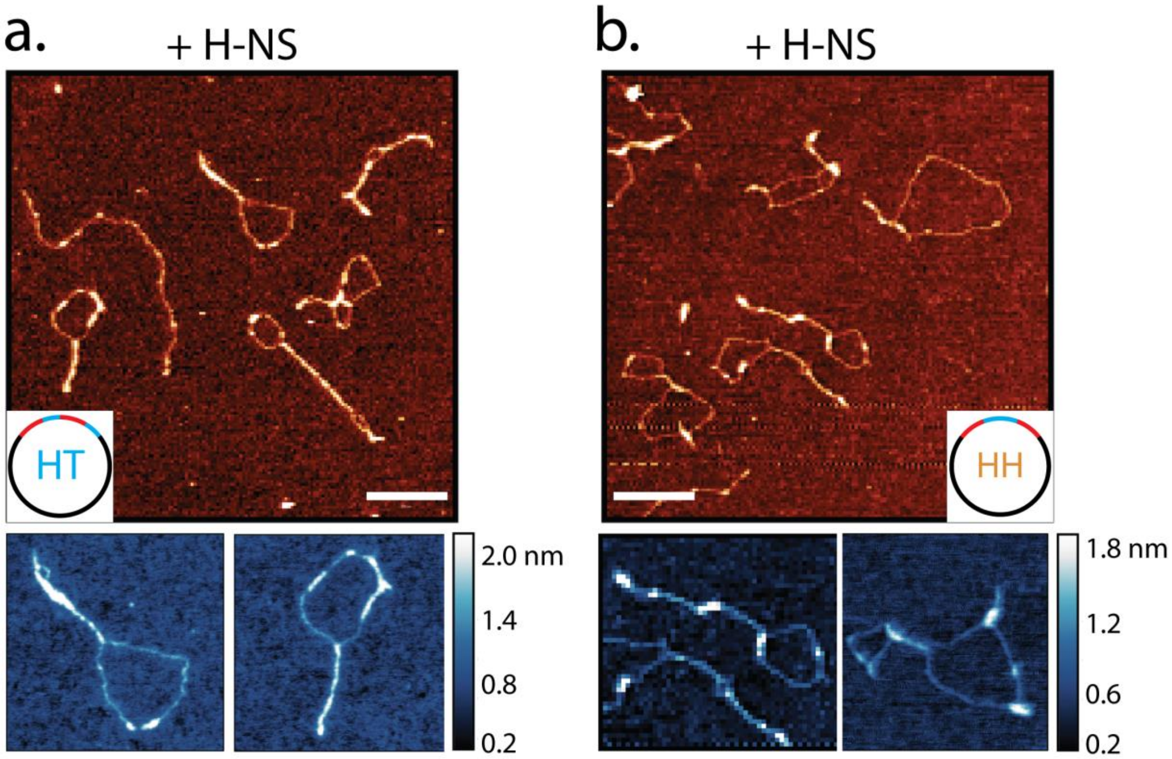


**Supplementary Figure 1.** AFM images of **a.** HT and **b**. HH circular constructs bound by H-NS protein. The binding of H-NS stabilizes DNA bridges. Scale bars 200nm.

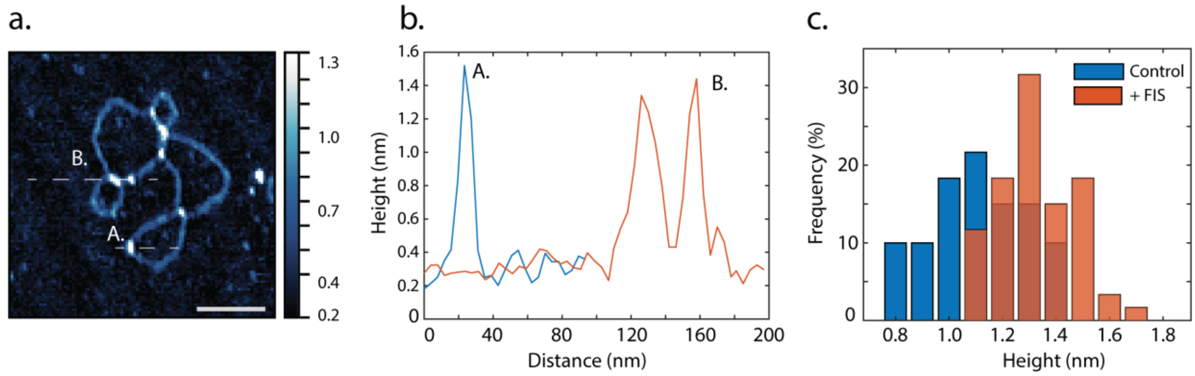


**Supplementary Figure 2. a**. AFM images of HT plasmid bound by FIS protein. Scale bar 100nm. **b**. Height measurement along the FIS proteins bound to the DNA marked as two dotted lines in panel a. **c.** Height distributions of DNA crossings in HT control (blue) and HT + FIS (red) samples. N=60 for both samples.

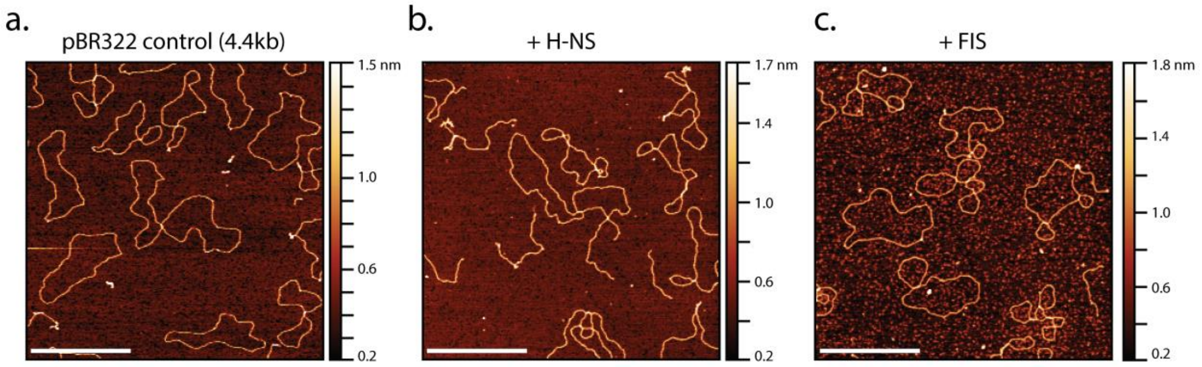


**Supplementary Figure 3.** AFM images **a.** Control pBR322 plasmid; **b.** H-NS bound pBR322 and **c.** FIS bound pBR322. Control plasmid is shows no or single DA crossings, H-NS bound plasmid shows bridged DNA regions, while FIS bound plasmid displays multiple DNA crossings. Scale bar 500nm.

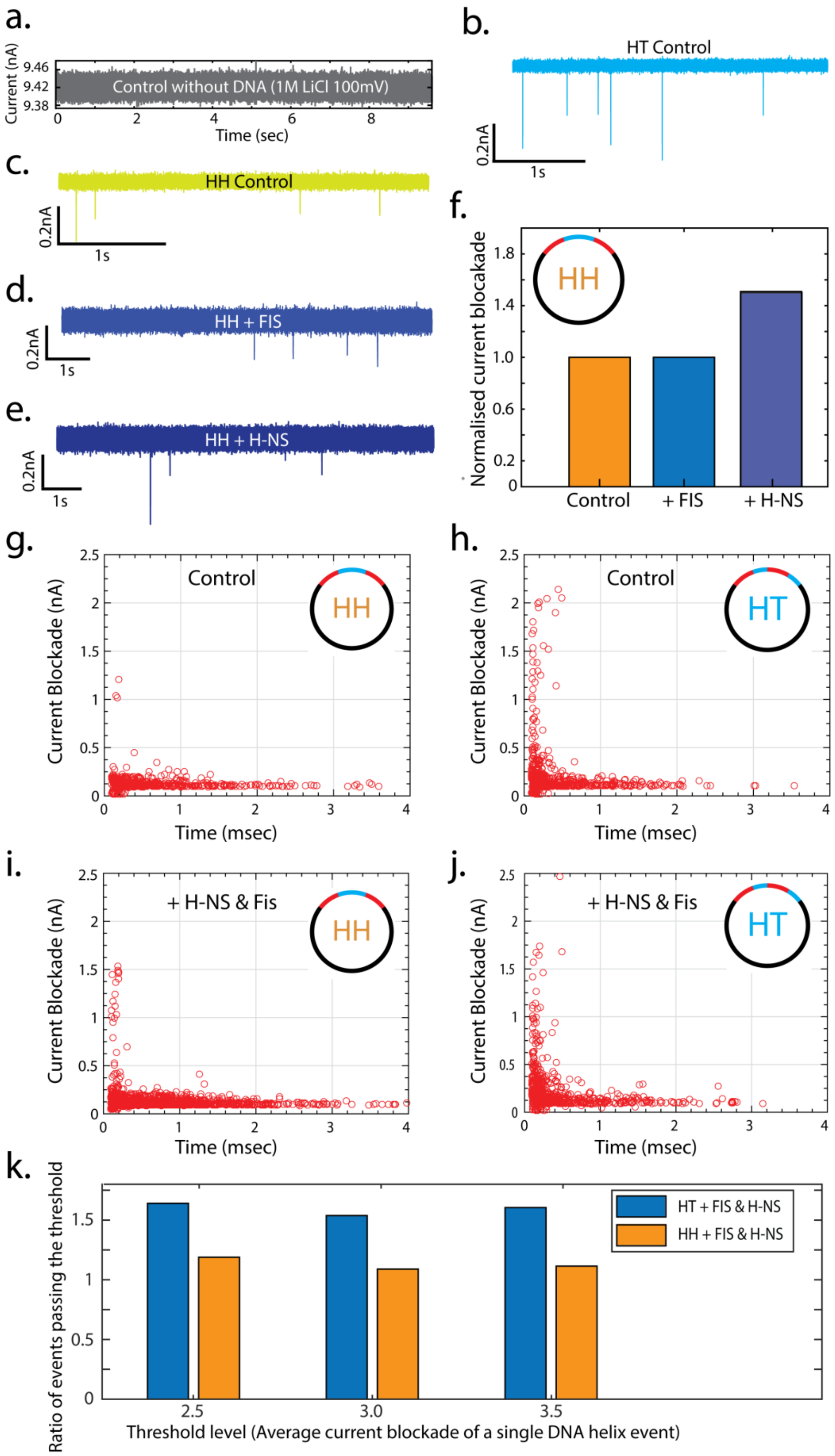


**Supplementary Figure 4.** Typical current traces of **a.** control sample without any DNA, not showing any translocations. **b.** HT control, **c.** HH control, **d.** HH construct bound with FIS, **e.** HH construct bound by H-NS, showing that with individual proteins we get deep and heterogeneous blockades, indicating various levels of nucleoprotein compaction. **f.** Normalized current blockade events for HH construct in the presence of single proteins. **g.** Current blockade vs. blockade time scatter plots for control (N=491) and **h.** H-NS & FIS bound HH samples (N=1812). **i.** Current blockade vs. blockade duration scatter plots for control (N=707) and **j.** H-NS & FIS bound HT samples (N=737). **k**. The ratio of events passing the current threshold for HT and HH construct in the presence of FIS and H-NS proteins (threshold is in units of average current blockade of a single DNA).

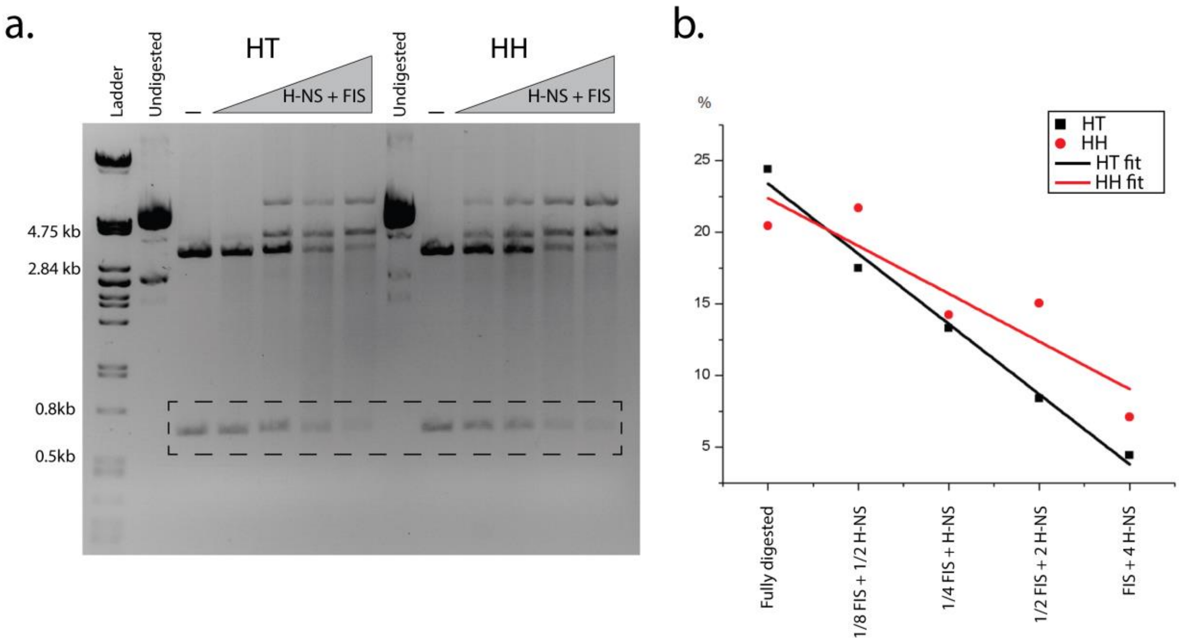


**Supplementary Figure 5**. Probing FIS and H-NS nucleoprotein complexes with BamH1 restriction endonuclease. **a.** Agarose gel electrophoresis. **b.** The change of the relative intensity of specific restriction bands (indicated with dotted rectangle on the panel a.) in each digestion reaction.

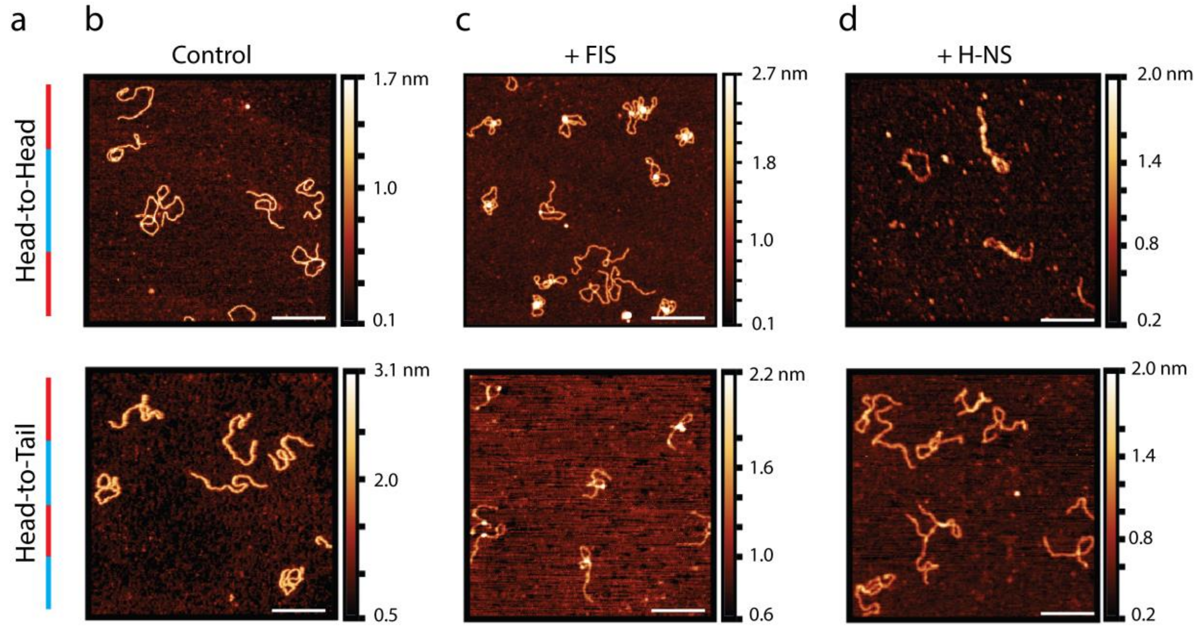


**Supplementary Figure 6. a.** Schematic of the sequence organisation of the 1.4kb HH and HT linear fragments. Large scale AFM images of **b.** control; **c.** FIS-bound and **d.** H-NS-bound linear fragments. Scale bar 200nm.

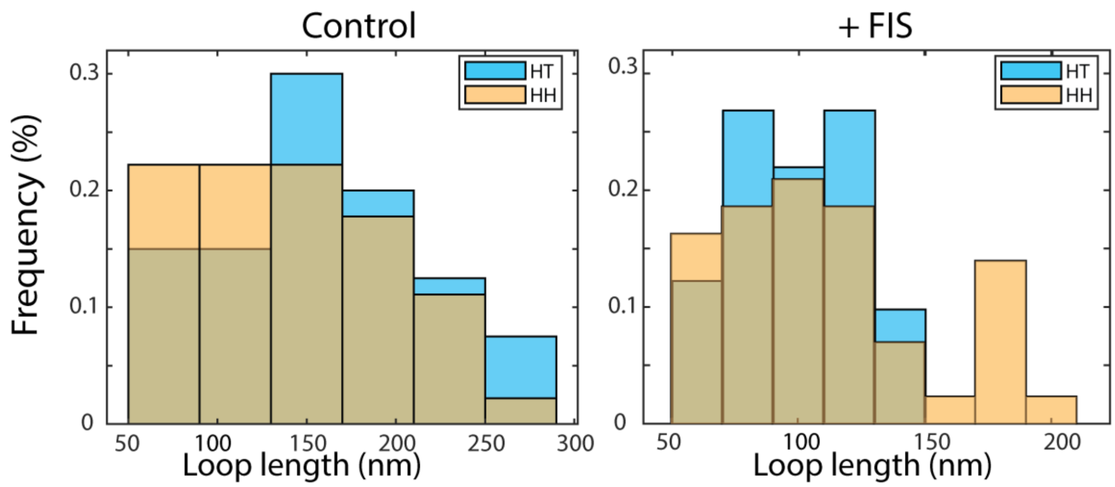


**Supplementary Figure 7.** Loop sizes formed by the binding of FIS on linear HT (blue bars) and HH (yellow bars) fragments.

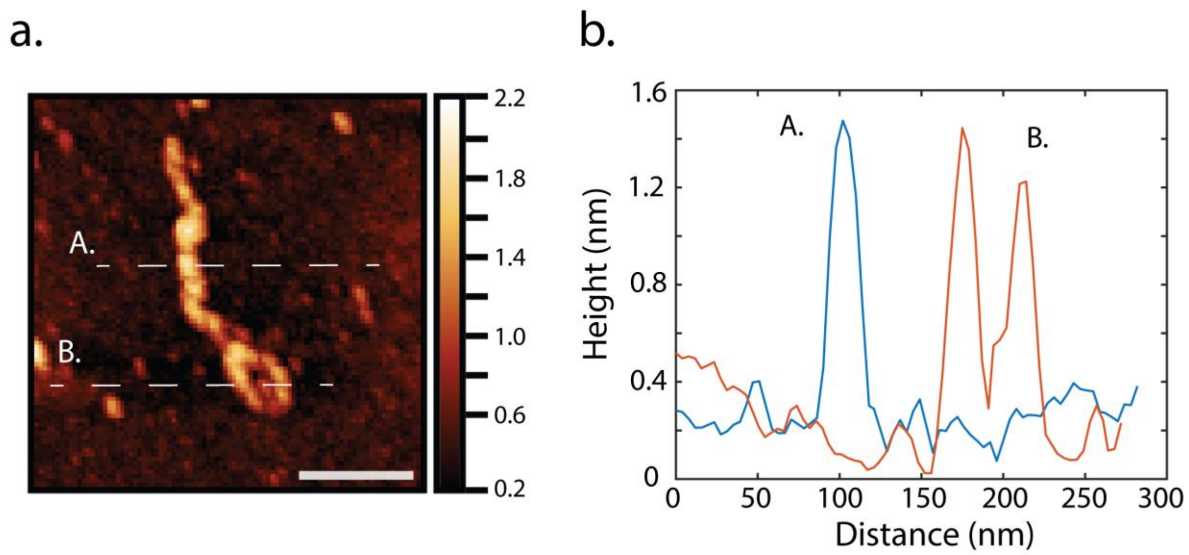


**Supplementary Figure 8. a**. AFM images of HH linear construct bound by H-NS protein. Scale bar 100nm. **b.** Height measurement along the H-NS proteins bound to the DNA marked as two dotted lines in panel a. The height profiles show that H-NS bridges two DNA duplexes (height profile A) as well as binds to single one inside loops (height profile B).

Supplementary Table 1. Shape parameters of linear nucleoprotein complexes

| Linear fragment DNA shape parameters | | | | | |
|---|---|---|---|---|---|
| Sample | Number | $L_{(contour)}$ [nm] | $\Delta L_{(contour)}$ | $l_p$ [nm] | $R_g$ [nm] |
| Head-Head (HH) | | | | | |
| Control | 110 | 445 ± 25 | X | 35 ± 5nm | 54 ± 13nm |
| +FIS : kb = 2.6 | 40 | 420 ± 30 | -6% | 27 ± 5nm | 42 ± 12nm |
| +H-NS : kb = 1.06 | 38 | 430 ± 30 | -3% | 43 ± 5nm | 52 ± 10nm |
| +FIS : H-NS : kb = 2.6 : 3.2 : 1 | 40 | 390 ± 50 | -12% | 50 ± 5nm | 85 ± 15nm |
| Head-Tail (HT) | | | | | |
| Control | 90 | 445 ± 30 | X | 40 ± 5nm | 58 ± 15nm |
| +FIS : kb = 2.6 | 35 | 380 ± 45 | -15% | 27 ± 5nm | 51 ± 12nm |
| +H-NS : kb = 1.06 | 45 | 430 ± 30 | -3% | 40 ± 5nm | 55 ± 11nm |
| +FIS : H-NS : kb = 2.6 : 3.2 : 1 | 35 | 460 ± 45 | +3% | 46 ± 5nm | 85 ± 18nm |
